# Priming of leaf litter decomposition by algae seems of minor importance in natural streams during autumn

**DOI:** 10.1101/353938

**Authors:** Arturo Elosegi, Angie Nicolás, John S. Richardson

## Abstract

Allochthonous detritus from terrestrial origin is one of the main energy sources in forested headwater streams, but its poor nutritional quality makes it difficult to use by heterotrophs. It has been suggested that algae growing on this detritus can enhance its nutritional quality and promote decomposition. So far, most evidence of this “priming” effect is derived from laboratory or mesocosm experiments, and it is unclear what its importance is under natural conditions. We measured accrual of algae, phosphorus uptake capacity, and decomposition of poplar leaves in autumn in open- and closed-canopy reaches in 3 forest and 3 agricultural streams. Chlorophyll a abundance did not change significantly neither with stream type nor with canopy cover, although some between open and closed reaches, although in some agricultural streams it was higher in open than in closed canopy reaches. Canopy cover did not affect either phosphate uptake capacity or microbial decomposition. On the other hand, although there was no effect of canopy cover on invertebrate fragmentation rate, a significant interaction between canopy cover and stream suggests priming occurs at least in some streams. Overall, the results point to a weak effect of algae on litter decomposition in natural streams during autumn.

## Introduction

Organic detritus from terrestrial origin, and leaf litter in particular, is one of the main energy sources for food webs in forested headwater streams (Fisher & Likens, 1973; Gessner *et al.*, 2010), where riparian shading limits primary production (Hill, Ryon & Schilling, 1995). This detritus tends to be dominated by recalcitrant compounds such as lignin and cellulose, is stoichiometrically imbalanced for the needs of consumers (Anderson, 1976; Hanson *et al.*, 1985; Lemoine, Giery & Burkepile, 2014; Evans-white & Halvorson, 2017), and thus, has low nutritional value (Brett *et al.*, 2017). Once in the water, leaf litter is colonised and “conditioned” by microbes, especially bacteria and aquatic hyphomycetes (Krauss *et al.*, 2011), whose lower C:N ratio enhances its overall palatability and nutritional value for invertebrate shredders (Hieber & Gessner, 2002; Findlay, 2010), which thus choose the most nutritious leaf patches (Fuller, Evans-White & Entrekin, 2015).

Yet, the lack of components such as essential fatty acids can still limit the nutritional value of leaves even after microbial conditioning (Torres-Ruiz & Wehr, 2010; Guo *et al.*, 2017). Recently, Lovatt *et al.* (2014) reported that leaf-litter leachate and light interact to promote algal biofilm accrual, which can in turn enhance the nutritional quality of leaf litter and thus promote the consumption by detritivore invertebrates (Guo *et al.*, 2016). This is an example of the *priming* effect, i.e., the enhancement of decomposition of recalcitrant organic matter by addition of labile carbon, which has been extensively documented in terrestrial ecosystems (Kuzyakov, 2010), much less frequently in aquatic ecosystems (Guenet *et al.*, 2010; Kuehn *et al.*, 2014). In the case of freshwater ecosystems, priming can involve the transfer of algal-produced C to heterotrophs (Kuehn *et al.*, 2014), or the direct consumption of algae by detritivore invertebrates (Ledger & Winterbourn, 2000; Franken *et al.*, 2005; Guo *et al.*, 2016), which would thus grow faster and have a larger effect on litter breakdown. It has even been suggested that consumers would preferentially use low-quality terrestrial carbon sources (e.g. leaf litter) for respiration, whereas they would preferentially use algae for secondary production (Karlsson, 2007).

Therefore, the priming effect could have important consequences for food webs, as well as for the management of stream ecosystems. Yet, most of the evidence on the priming effect is derived from experiments either in the laboratory or in artificial channels. For instance, Danger *et al.* (2013) in a factorial mesocosm experiment with light and nutrients, showed that under low nutrient concentrations diatom growth enhanced litter decomposition. The importance of the priming effect under field conditions is currently unclear. In theory, priming should be most important in periods of stable baseflow and high light-availability, which promote the accrual of algal biomass (Biggs, 1995). However, it is possible that priming could be less important in high-flow periods, in strongly shaded reaches or in periods of large accumulations of detritus, as heterotrophic microbes have been shown to outcompete algae for limiting nutrients (Mindl *et al.*, 2005; Danger, Benest & Lacroix, 2007). In particular, in many temperate streams the bulk of leaf fall occurs in autumn (Abelho, 2001), a period in which the differences in light availability between open and closed reaches decrease as a consequence of leaf fall and shorter daylight period, which might reduce the real effect of priming. Also, it has been suggested that the priming effect is more important in nutrient-poor conditions, where algae would release more labile C exudates (Guenet *et al.*, 2010), but there is so far little empirical evidence of this.

The objective of the present study was to test the importance of priming in natural streams during the peak of litterfall. More specifically, we assessed the effects of autumnal riparian cover on biofilm accrual and activity, and on litter decomposition in streams of contrasting nutrient status. Our hypotheses were: i) that biofilm accrual will be higher at open than at closed reaches, ii) that higher biofilm accrual will boost litter breakdown at nutrient-poor but not at nutrient-rich streams, and iii) that the effect will be higher for invertebrate fragmentation than for microbial decomposition.

## Methods

### Study sites

The experiment was performed in six streams, 3 forested (F1 to F3) and 3 agricultural (A1 to A3), near Vancouver, British Columbia, Canada (Table 1). The forest streams are located in the Malcolm Knapp Research Forest (Maple Ridge, BC), a 52 km^2^ area owned by the University of British Columbia. It is almost entirely covered by forest, dominated by Douglas-fir (*Pseudotsuga menziesii*), western hemlock (*Tsuga heterophylla*) and western red cedar (*Thuja plicata*), and with red alder (*Alnus rubra*) and black cottonwood (*Populus trichocarpa*) abundant in the riparian areas. The lithology is granitic, soils acidic and streams in the area have extremely low conductivity and nutrient concentrations. The agricultural streams are located ca. 30 km further south, in the Fraser River Delta (Surrey and Aldergrove, BC), a region with intense agricultural activities, including berry farms and dairies, and scattered urban areas. The same tree species occur, but the forest cover is much lower than in the forest streams, grasses and bushes becoming more important. Lithology is dominated by glacial till, streams are circumneutral and richer in nutrients (Shupe, 2017).

**Table 1.** Main characteristics of the studied streams. Land cover data obtained from the Land Use 2010 map by the Canada Government https://open.canada.ca/data/en/).

| Stream | Name | Coordinates | Stream<br>order | Basin<br>(Km<sup>2</sup>) | Forest<br>(%) | Agriculture<br>(%) | Urban<br>(%) | Water<br>(%) |
| --- | --- | --- | --- | --- | --- | --- | --- | --- |
| F1 | Blaney Creek | 49°16'14" N<br>122°35'20" W | 2 | 8.74 | 94 | 0 | 0 | 6 |
| F2 | Spring Creek | 49°16'15" N<br>122°34'30" W | 2 | 2.55 | 100 | 0 | 0 | 0 |
| F3 | Mayfly Creek | 49°15'06" N<br>122°32'51" W | 2 | 1.14 | 100 | 0 | 0 | 0 |
| A1 | Pepin Creek | 49°00'14" N<br>122°28'18" W | 2 | 15.38 | 33 | 29 | 35 | 0 |
| A2 | Bertrand Creek | 49°01'58" N | 3 | 30.87 | 27 | 34 | 35 | 0 |
|  |  | 122°32'04" W |  |  |  |  |  |  |
| A3 | Little Campbell | 49°01'60" N | 3 | 28.74 | 56 | 21 | 21 | 2 |
|  | River | 122°41'06" W |  |  |  |  |  |  |

The climate in the Vancouver region is moderate oceanic, with an annual mean temperature near 11 °C and a precipitation ranging from 1500 mm per year in the coast to over 3000 mm in the mountains. Summers are drier, and autumn tends to be very rainy.

In each stream we selected two nearby (less than 200 m apart) riffle reaches, one with open canopy (O), and the other with closed canopy (C).

### Biofilm

To measure biofilm accrual and activity we used *biofilm carriers*. These are standard materials where biofilm can attach, designed to promote self-purification in aquaria by encouraging growth of bacteria inside the water pump. We used Biofilter balls (Marineland, United Pet Group, Spectrum Brand Inc., Blacksburg, VA), which consist of hollow plastic spheres, 2 cm in diameter, with a surface made of thin plates, the space between plates allowing water flow as to promote biofilm growth. Two such balls were tied with fishing line to each of the rebars used to deploy litterbags (see below), at mid depth. They were recovered together with the litterbags, enclosed by pairs in 50 mL Falcon tubes with filtered stream water, and carried to the laboratory in an ice chest to perform a phosphate-uptake bioassay.

Once in the laboratory, the stream water was replaced by an acclimation solution (1 L of Perrier water in 4 L of distilled water), designed to ensure a sufficient supply of micronutrients, and the Falcons were incubated for 30 min in a LabRoller rotator at minimum speed inside a Cenviron environmental chamber at 8 °C temperature and 50 μmol of PAR (Apogee quantum sensor SQ100). After this acclimation period, the water in the Falcons was replaced by a 5 μM solution of PO_4_^3-^, prepared by dissolving H_2_NaO_4_P·H_2_O in the acclimation solution, and were incubated for 1 h. An additional set of five pairs of uncolonized biofilm carriers was used as blanks. After this period, water was filtered from the Falcons (Whatman GF/F) and frozen to later analyse the remaining P concentration (see below), and the biofilm carriers were frozen in the Falcons. The uptake rate of phosphorus (*Up*, in mg P h^−1^) was calculated as:

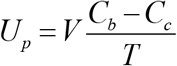

where *V* is the volume of incubation solution (L), *C*_*b*_ is the final concentration of P (mg P L^−1^) in the bank treatment (uncolonized biofilm carriers), *C*_*c*_ is the final P concentration in the colonized treatment and *T* is the incubation time (h).

Biofilm carriers were later thawed, kept by pairs in 20 mL of acetone overnight at 4 °C, and their content of chlorophyll was measured fluorometrically (TD-700, Turner Designs), using uncolonized biofilm carriers in acetone as a blank.

### Leaf breakdown

We studied the breakdown of black cottonwood leaves, a common riparian tree in the area with relatively poor-quality leaves (Crutsinger *et al.*, 2014), which, thus, could be more prone to priming. Recently fallen leaves were collected below five adjacent trees in Burnaby (BC, Canada), taken to the laboratory and air-dried. Batches of 3.0 (±0.1) g of dry leaves were enclosed in coarse (9 mm) or fine (250 μm) mesh bags. One fine bag was enclosed inside each coarse bag, thus ensuring that they were subject to the same environmental conditions. Bags were taken to the field and tied with cable binders simultaneously to the biofilm carriers, to five metal rebars per reach (1 double bag per rebar), anchored in riffle sections. Bags were retrieved after 2 months of incubation, enclosed in zip-lock bags and carried to the laboratory in an ice chest. There, they were opened, the remaining leaf material was cleaned with tap water on a 250-μm sieve, and the ash-free dry mass (AFDM) was measured gravimetrically after drying (60 °C, 96 h) and ashing (500 °C, 4 h). The initial ash-free dry mass was calculated from 10 additional batches of dry leaves, which were dried and ashed as explained. Breakdown rates were calculated according to the negative exponential model (Petersen & Cummins, 1974) with time in degree-days. Following Lecerf (2017) we calculated the mean litter fragmentation rate (*λ*_*F*_) as

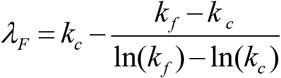

where *λ_F_* is the mean fragmentation rate and *k_c_* and *k_f_* the breakdown rates in coarse and fine mesh bags, respectively.

### Environmental variables

On 5 occasions during the decomposition experiment, we measured wetted cross-section with a ruler and a measuring tape, and water velocity (current meter Swoffer 2100) to calculate stream discharge. Additionally, we measured water temperature, conductivity and pH (field probe YSI Pro1030) and dissolved oxygen (field probe YSI ProODO), and collected water samples, which were filtered (Whatman GF/F), carried to the laboratory in an ice chest, and frozen immediately for analysis of nutrients. The concentrations of nitrate, nitrite, ammonium and phosphate in water were determined by an OI-Analytical “Alpkem Flow System IV” automated chemistry analyser at the Department of Analytical Chemistry, British Columbia Ministry of Environment and Climate Change Strategy, Victoria, British Columbia.

Additionally, water temperature was recorded continuously at each reach by Onset TidBit v2 temperature loggers tied to one of the rebars. Riparian cover was estimated from zenithal photos taken with a 28 mm (35 mm equivalent) lens (camera Fujifilm X20), which were later processed to maximize contrast and then analysed with ImageJ, a free software downloadable from https://imagej.nih.gov/ij/ to measure total image brightness, which was converted into cover.

### Statistical analyses

We used linear mixed effect models to contrast stream water characteristics (IBM SPSS V.24) with stream type (agricultural vs forested) and canopy cover (open vs closed) as fixed factors, and stream identity as random effect with five levels. Nutrient variables (PO_4_-P and TIN) were log transformed to meet normality assumptions. Significance for each term was assessed by model comparisons using likelihood ratio test (Crawley, 2007).

## Results

The experiment started in late September 2017, after an unusual dry summer that caused extremely low baseflows in the streams in the area. The weather suddenly changed after the 16^th^ of October, when large storms affected the area for 5 days (http://climate.weather.gc.ca/). Floods scoured all bags and most of the biofilm carriers deployed in Blaney Creek (stream F1), which led us to remove this stream from the study. After these floods, weather remained fairly dry from 26^th^ October to 7^th^ November, and then it rained almost every day until the end of the experiment, although no large floods were registered.

Riparian cover at the beginning of the experiment ranged from 0 to 45% at the open reaches, whereas it was higher than 76% at the closed reaches (Table 2). It decreased along the experiment in all reaches, as deciduous trees lost their leaves, but the closed reaches still maintained riparian cover higher than 55%. Average discharge over the study period ranged from 129 L s^−1^ in stream A1 to 586 Ls^−1^ in stream A2. The average temperature was higher in agricultural than in forest streams (9.2-9.9 vs 7.0-8.5 °C, respectively), as were pH (7.0-7.4 vs 6.4-6.5) and conductivity (163-252 vs 19 μScm^−1^). On the contrary, oxygen concentration tended to be higher at forest (11.0-11.9 mgL^−1^) than at agricultural (9.3-10.1 mgL^−1^) streams (Table 2). Average TIN concentrations were below 0.30 mg N L^−1^ in forest streams, whereas they ranged from 0.47 to 1.81 in agricultural streams. SRP concentrations were below 6 μg P L^−1^ in forest streams, whereas they ranged from 12.8 to 35.0 in agricultural streams. Linear mixed models detected statistically significant difference between agricultural and forest streams for oxygen concentration (F_1,4_ = 64.6; p=0.001), pH (F_1,4_ = 58.5; p=0.002), conductivity (F_1,4_ = 239.2; p<0.001), TIN (F_1,4_ = 16.1; p=0.016) and phosphate (F_1,4_ = 27.8; p=0.006), whereas differences were marginal for temperature (F_1,4_ = 5.1; p=0.088). None of these variables showed statistically significant differences between open and closed reaches.

**Table 2.**
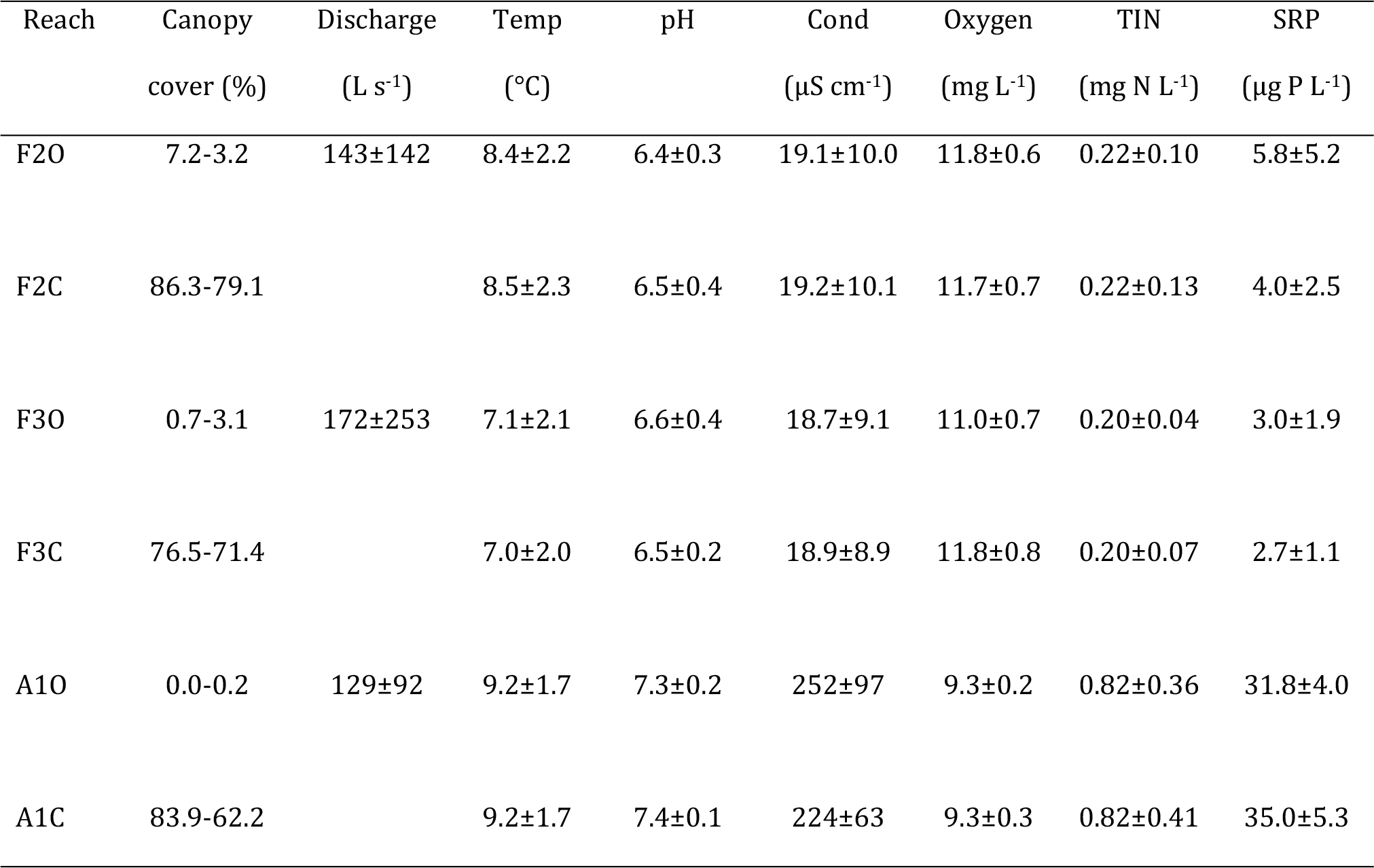
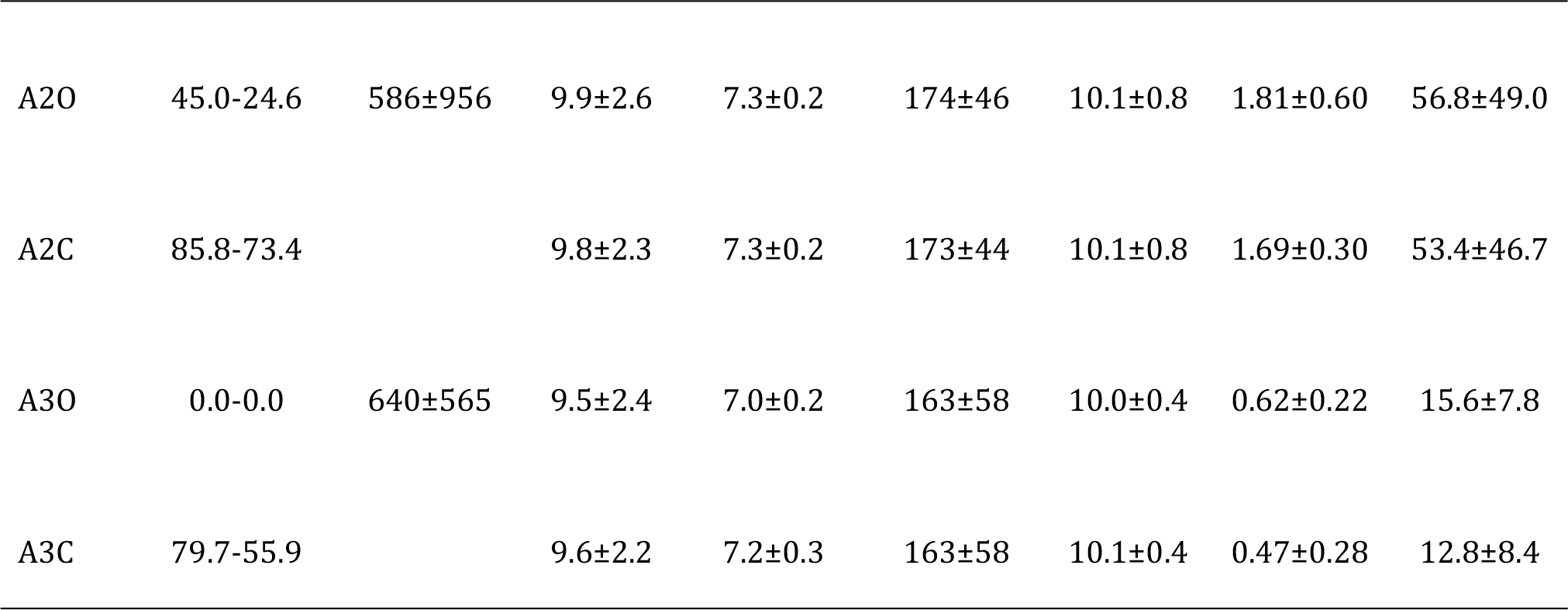
Environmental characteristics of the study reaches. F stands for forest, A for agricultural, O for Open, C for Closed. Canopy cover values are point values measured at the beginning and at the end of the experiment. The rest of the values are mean ± SD of 5 measurements during the experiment. TIN (total inorganic nitrogen) is the sum of nitrate, nitrite and ammonium. SRP, soluble reactive phosphorus. Discharge was only measured in one site per stream, where it was most convenient, not necessarily in any of the sampling reaches.

| Reach | Canopy<br>cover (%) | Discharge<br>(L s<sup>-1</sup>) | Temp<br>(°C) | pH | Cond<br>(μS cm<sup>-1</sup>) | Oxygen<br>(mg L<sup>-1</sup>) | TIN<br>(mg N L<sup>-1</sup>) | SRP<br>(μg P L<sup>-1</sup>) |
| --- | --- | --- | --- | --- | --- | --- | --- | --- |
| F2O | 7.2-3.2 | 143±142 | 8.4±2.2 | 6.4±0.3 | 19.1±10.0 | 11.8±0.6 | 0.22±0.10 | 5.8±5.2 |
| F2C | 86.3-79.1 | | 8.5±2.3 | 6.5±0.4 | 19.2±10.1 | 11.7±0.7 | 0.22±0.13 | 4.0±2.5 |
| F3O | 0.7-3.1 | 172±253 | 7.1±2.1 | 6.6±0.4 | 18.7±9.1 | 11.0±0.7 | 0.20±0.04 | 3.0±1.9 |
| F3C | 76.5-71.4 | | 7.0±2.0 | 6.5±0.2 | 18.9±8.9 | 11.8±0.8 | 0.20±0.07 | 2.7±1.1 |
| A1O | 0.0-0.2 | 129±92 | 9.2±1.7 | 7.3±0.2 | 252±97 | 9.3±0.2 | 0.82±0.36 | 31.8±4.0 |
| A1C | 83.9-62.2 | | 9.2±1.7 | 7.4±0.1 | 224±63 | 9.3±0.3 | 0.82±0.41 | 35.0±5.3 |
| A2O | 45.0-24.6 | 586±956 | 9.9±2.6 | 7.3±0.2 | 174±46 | 10.1±0.8 | 1.81±0.60 | 56.8±49.0 |
| A2C | 85.8-73.4 |  | 9.8±2.3 | 7.3±0.2 | 173±44 | 10.1±0.8 | 1.69±0.30 | 53.4±46.7 |
| A3O | 0.0-0.0 | 640±565 | 9.5±2.4 | 7.0±0.2 | 163±58 | 10.0±0.4 | 0.62±0.22 | 15.6±7.8 |
| A3C | 79.7-55.9 |  | 9.6±2.2 | 7.2±0.3 | 163±58 | 10.1±0.4 | 0.47±0.28 | 12.8±8.4 |

Chlorophyll *a* per pair of biofilm carriers ranged from 0.83±0.43 μg (mean±SD) in F2 to 10.53±7.99 in A1 (Fig. 1). Differences were not significant between stream types (LMM, F_1,3_ = 2.4; p=0.216) nor between open and closed reaches (F_1,3_ = 3.5; p=0.154), and the interaction canopy*type was neither significant (F_1,3_ = 2.12; p=0.239). Nevertheless, there were statistically significant differences of the stream factor (F_3,3_ = 13.5; p=0.030) as well as for the canopy*stream interaction (F_3,37_ = 3.2; p=0.035), showing that the canopy effect was significant at some streams. In these cases, open reaches had more chlorophyll than closed ones.

**Figure 1.**
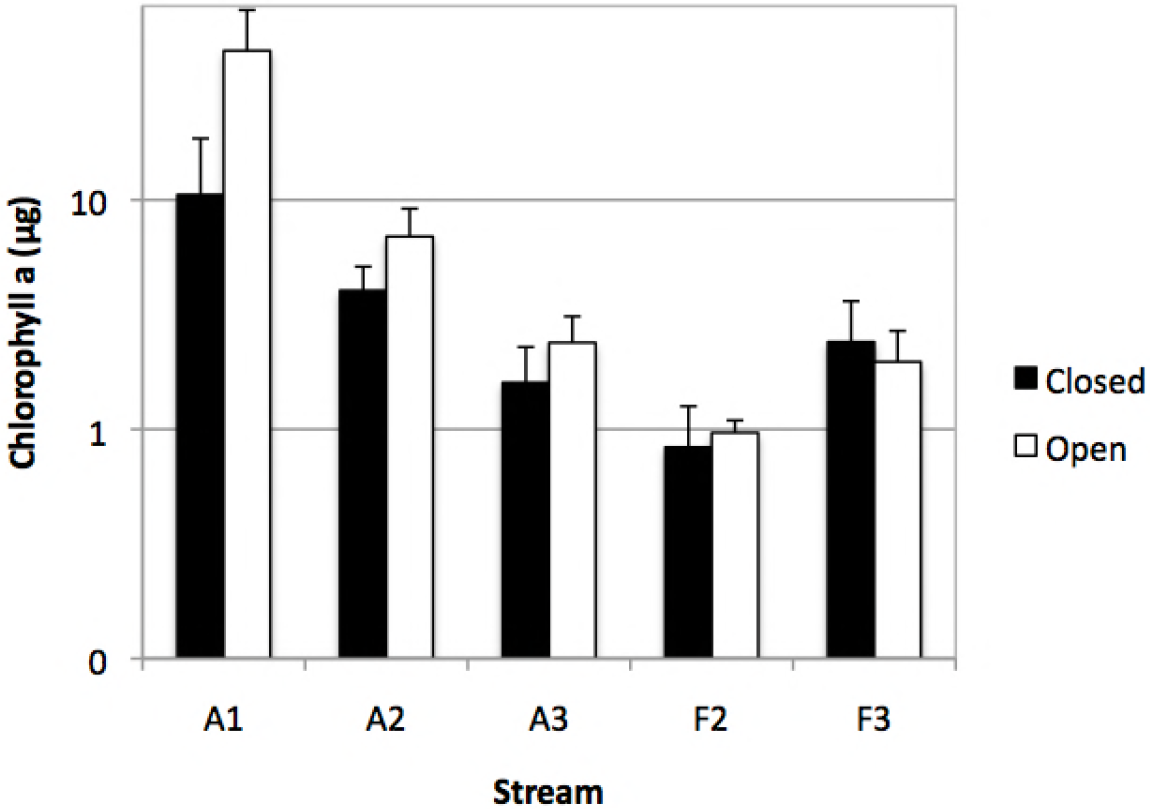
Chlorophyll *a* per pair of biofilm carriers in open and closed reaches of agricultural (A) and forest (F) streams. Note logarithmic scale.

Phosphate uptake in the bioassay performed with biofilm carriers ranged from 0.96±0.28 μg P h^−1^ in reach F2C to 1.52±0.37 in reach F3C (Fig. 2). The uptake per unit of chlorophyll ranged from 32.2±12.4 to 1582±382 μg P mg Chl^−1^ h^−1^ (Fig. 2). It was lowest in streams A1 and A2, the two ones with highest basal concentrations of SRP in water. None of the factors and interactions tested by LMM resulted statistically significant for phosphate uptake (p>0.05 in all cases), but there were significant differences in uptake per unit of chlorophyll for the factor stream (F_3,3_ = 10.3; p=0.044) and the interaction canopy*stream (F_3,37_ = 4.0; p=0.014).

**Figure 2.**
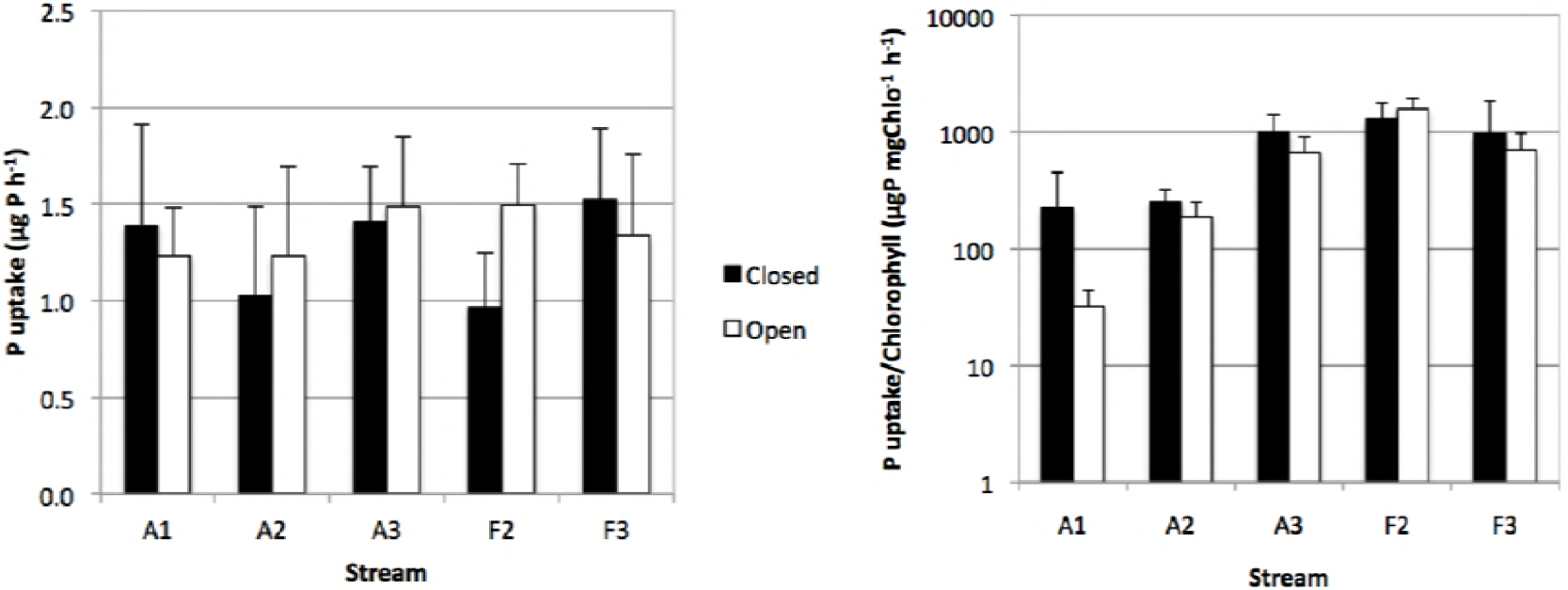
Left. Phosphate uptake by biofilm carriers in open and closed reaches of agricultural (A) and forest (F) streams. Right. Efficiency in phosphate uptake of chlorophyll in biofilm carriers.

After 2 months of incubation, the leaf AFDM remaining in fine mesh bags ranged from 18.02±6.26 in reach AO3 to 49.44±2.51% in FC2 (Fig. S1). No clear differences could be detected either between agricultural and forest streams, nor between closed and open reaches. The leaf AFDM remaining in coarse mesh bags ranged from 1.80±1.96 in AO3 to 47.17±2.32 in FO3. It was higher in forest than in agricultural streams, but no obvious differences were detected between open and closed reaches. The temperature-corrected breakdown rates ranged from 0.0015±0.0001 to 0.0032±0.0006 dd^−1^ in fine mesh bags and from 0.0018±0.0003 to 0.0082±0.0021 dd^−1^ in coarse mesh bags. In fine mesh bags there were no obvious differences between agricultural and forest streams, but in coarse mesh bags agricultural streams tended to have higher breakdown rates (Fig. 3). No clear differences were seen between open and closed reaches. LMM showed the factors stream type and canopy cover and their interaction to have no statistically significant effect on fine-mesh decomposition rate, but stream had (F_3,3_ = 33.8; p=0.008). For coarse mesh bags the only statistically significant difference was attributed to the canopy*stream interaction (F_3,36_ = 3.45; p=0.027), although the direction of the difference was not consistent among streams.

**Figure 3.**
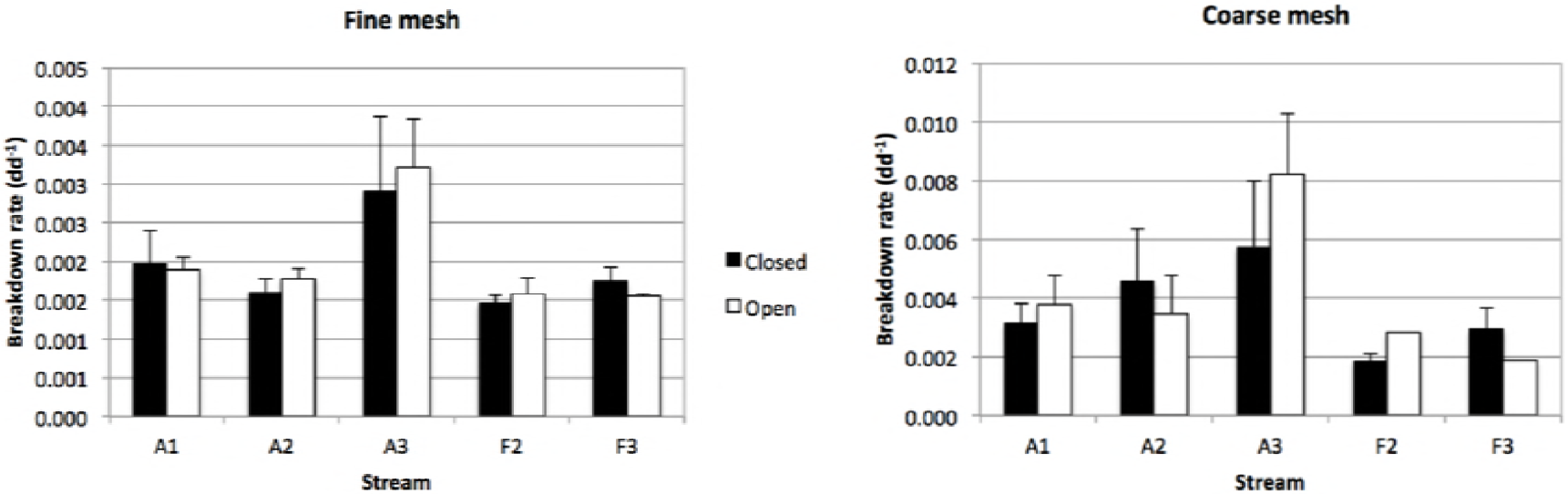
Temperature-corrected breakdown rates in fine (left) and coarse (right) mesh bags in open and closed reaches of agricultural (A) and forest (F) streams. Note the different scale of Y axes.

Finally, the fragmentation rate, i.e., the contribution of shredders and physical abrasion to total breakdown, ranged from 0.0011 to 0.0259 d^−1^, and tended to be higher in agricultural than in forest streams (Fig. 4), although differences between stream types were not statistically significant. LMM only found a statistically significant effect for the interaction between canopy cover and stream (F_3,36_ = 4.83; p<0.006), thus showing that fragmentation rate was higher in open than in closed reaches of some streams.

**Figure 4.**
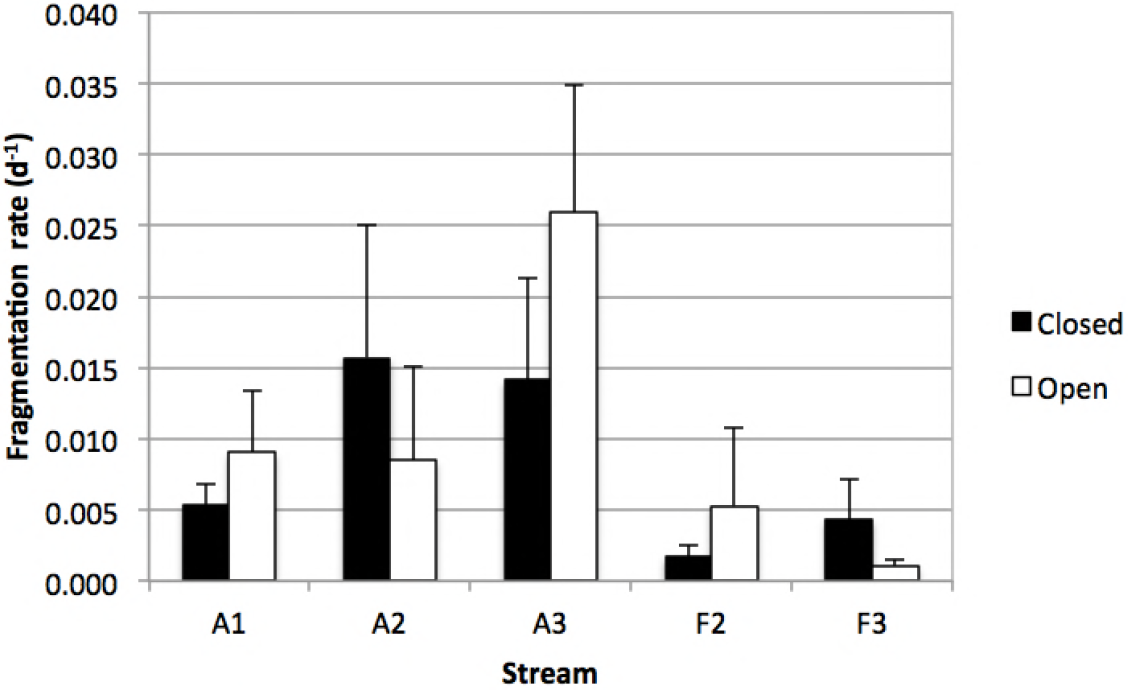
Fragmentation rates in coarse mesh bags in open and closed reaches of agricultural (A) and forest (F) streams.

## Discussion

Our results point to a weak priming effect by algae on autumn litter decomposition in the study streams. In agricultural streams there was a trend for open reaches to have more chlorophyll than closed ones, but the differences were not consistent and did not translate into faster litter decomposition. Although stream*canopy interactions showed differences between open and closed reaches in some streams, the direction of the difference was not consistent among streams. Thus, our overall results suggest that other environmental factors override the potential effect of algal growth on leaf litter dynamics.

Our experimental design, with open and closed reaches very close to each other, aimed at reducing the potentially confounding effects of other environmental variables apart from canopy cover. Among our forest streams open reaches are rare and very short, whereas among our agricultural streams usually there is an alternation of short open and closed reaches. Therefore, none of the environmental variables measured in addition to canopy cover showed systematic differences among reach types, as water quality variables usually respond to riparian cover at longer scales, on the order of hundreds to thousands of metres (Allan, 2004; Johnson, 2004). On the other hand, the differences in canopy cover between open and closed reaches decreased but did not disappear over the study period, and therefore, the changes found among reach types can be attributed to canopy cover.

Under these conditions, algal biomass showed no response to canopy cover in the forest streams, thus suggesting algal growth there to be limited by factors other than light, probably nutrients. The concentration of phosphorus in our forest streams was close, when not below, detection limits. Additionally, high N:P atomic ratio ranged from 52 (A1C) to 164 (F3C), suggesting P to be the limiting nutrient in all our sites (Elser *et al.*, 2000; Sterner *et al.*, 2008). As the winter approaches, other factors such as low water temperature, decreased day length, low elevation of the sun in the horizon, frequent cloudy days and high flow tend to limit algal growth (Izagirre & Elosegi, 2005), which probably explains the lack of differences in our forest, nutrient-limited streams. Long baseflow periods promote biofilm accrual (Ponsatí *et al.*, 2015), whereas floods scour algae (Francoeur & Biggs, 2006), and thus, in mountain streams frequent floods can override the effects of canopy cover on algal biomass (Boulêtreau *et al.*, 2008). It is likely that, even if there were no differences in chlorophyll content, the algal assemblages differed between open and closed reaches, as algal taxa differ in their competitive abilities depending on light and nutrient availability (Franken *et al.*, 2005; Litchman & Klausmeier, 2008; Lange, Townsend & Matthaei, 2016). However, even if this was the case, overall it had no measurable effect on our decomposition rates.

Contrary to our expectations, phosphate uptake capacity, a proxy for biofilm activity, did not differ between forest and agricultural streams, or between open and closed reaches. In general, the metabolic activity of stream biofilm depends on its biomass (Haggerty *et al.*, 2014), whereas the uptake per unit biomass tends to be greatest at oligotrophic, nutrient-poor reaches (Mulholland, 1996). Nevertheless, internal phosphorus recycling at the biofilm level gains importance in nutrient-rich reaches (Mulholland *et al.*, 1995), which usually reduces uptake rate (Proia, Romaní & Sabater, 2017) and results in our bioassay yielding highest uptake rate in moderately enriched streams (Arturo Elosegi, unpublished data). The small differences we found in P uptake rate per unit of chlorophyll suggest a part of the algal biomass to be not very active, probably as a consequence of the senescence of algal mats by the end of autumn, which reduces the biological activity even if the biomass is large (Izagirre *et al.*, 2008). Interestingly, the weak trends found between open and closed reaches point towards a higher nutrient uptake per unit biomass in the latter, which again, was contrary to our expectations, and could be related to site-specific characteristics such as flow velocity.

Microbial decomposition, as measured by temperature-corrected breakdown rate in fine-mesh bags, did not differ between agricultural and forest streams, or between open and closed reaches, although the random factor stream was statistically significant. This is a surprising result, as moderate nutrient concentrations as those found in our agricultural streams have been shown to promote litter breakdown (Gulis, Ferreira & Graca, 2006), although effects are much more clear for total decomposition (Chauvet *et al.*, 2016). For coarse mesh bags, although there seemed to be a trend for litter to decompose faster at agricultural streams, the differences in temperature-corrected breakdown rates were again not statistically significant, contrasting with clear patterns shown elsewhere (Woodward *et al.*, 2012). Nevertheless, the statistically significant interaction between canopy cover and stream suggested a weak priming effect in some of the streams. We can only speculate about the reasons for this difference occurring only in some streams, but it is likely that local factors affected differently the two reaches studied in one stream, thus overriding the weak effects of priming. Indeed, litter breakdown is sensitive to small differences in flow velocity (Dewson, James & Death, 2007), sediment deposition (Piggott *et al.*, 2012) or biological communities (Handa *et al.*, 2014), factors that can change at the micro-habitat scale (Elosegi, Flores & Díez, 2011). In our case, shredders were present in all sites (Arturo Elosegi, personal observation), including agricultural streams. Intense human activities can result in local extinction of large shredders such as crustacean gammarids, and thus impact litter breakdown (Hladyz *et al.*, 2011), but our streams seem not to have crossed this impairment threshold.

The trend towards higher fragmentation rates suggests that invertebrates were even more important in our agricultural than in our forest streams, which is usually not the case (Hladyz *et al.*, 2011). Again, the interaction between stream and canopy cover was statistically significant, suggesting that the priming effect can be important at least in some streams.

Overall, our results suggest that algal priming of litter decomposition, a key ecosystem process (Hagen *et al.*, 2012), is at best weak in natural streams during autumn, when the environmental conditions become less favourable for algal growth. The relevance of priming in streams remains a controversial effect. Although the nutritional quality of biofilms tends to be greater when algae are present (Huggins, Frenette & Arts, 2004) and much of the nitrogen assimilated by stream consumers is of algal origin (Brett *et al.*, 2017), the effects on litter decomposition are often unclear. Most of the research has been performed in the laboratory or in artificial streams, and even there results are far from unequivocal. For instance, a recent laboratory experiment (Guo *et al.*, 2016) found enhanced nutrients but not light to promote shredder biomass and breakdown rate, contrasting with other experiments (Franken *et al.*, 2005), who found no effect on shredder-mediated decomposition, although light promoted the growth of *Asellus* and *Gammarus* crustaceans. (Guo *et al.*, 2016) attributed the lack of effect to the small difference between their two light levels (21 and 114 μmol m^−2^ s^−1^). We did not measure differences in light levels between our open and closed reaches, but it is likely that they were rather small and decreased during the experiment, as a consequence of leaf fall, shortening of the day and increased cloudiness. As for field experiments, although some authors (Kuehn *et al.*, 2014) found exposure to light to stimulate decomposition in the field, their “shadow” treatment was artificial, as they covered the channel with cloth. Similarly, Vonk *et al.* (2016) found that, whereas in microcosms, invertebrates showed a preference for artificial substrate with poly-unsaturated fatty acids added, this effect could not be detected in the field, probably because it was overruled by unknown sources of variation. Priming could be more important in summer, when the differences in light availability between open and closed reaches are largest and the long baseflow periods make biological effects more marked.

## Acknowledgements

We are grateful to the landowners and managers of the nearby lands for granting access to the study sites, to the members of Richardson Laboratory (UBC) for assisting in the field and in the laboratory, to the staff of the Forestry Faculty (UBC) for technical support, and to Aitor Bastarrika (UPV/EHU) for quantifying land cover. The project has been funded by the Basque Government (IT951-16) and the Natural Sciences and Engineering Research Council (Canada). Arturo Elosegi enjoyed a grant from the Basque Government for a research stay at the University of British Columbia.

